# Alternative splicing in 300 human cells and tissues is sparsely distributed and strongly associated with cell identity

**DOI:** 10.1101/2025.01.06.631625

**Authors:** Angelita Liang, Marc R Wilkins

## Abstract

In eukaryotes, alternative splicing allows RNA sequences to be changed as required in their cellular context. While the use of gene expression without considering transcript isoforms is commonly used to classify cell types, the significance of alternative splicing to cell identity is not fully understood. Using data-driven classification and clustering methods, we analysed the RNA Atlas, an ultra-deep total RNA and polyA capture sequencing dataset obtained from 300 different types of cell and tissue samples across the human body. We show that splice inclusion values can distinguish cell identity effectively, and sometimes better than gene expression. These specific differences in alternative splicing are biologically relevant, with many splice events expressed in a limited number of cell types. We show that distributions of splice inclusion levels are “sparser” than gene expression levels, being discontinuous and containing outliers – akin to a control dial with a few settings. Gene expression levels display finer variation, taking on a diverse range of values akin to a control dial with many increments. Complementary analysis of comparable RNA-Seq samples from the ENCODE dataset yielded consistent results regardless of the splice event or gene expression quantification tools used, as well as the choice of RiboMinus^TM^ or polyA capture-derived RNA libraries. Our results highlight the utility of alternative splicing data in data-driven and functional analyses, and the unique relationship between alternative splicing and cell identity.

## INTRODUCTION

In eukaryotes, alternative splicing of RNA is an essential process that enables the transcription of diverse RNA sequences from a single gene by assembling exons in a configuration appropriate for their cellular context (Chen and Manley, 2009). This process is subject to tight upstream regulation by splicing factors which often function in a combinatorial manner (Dvinge, 2018; Matlin et al., 2005). Transcript isoforms arising from alternative splicing participate in a vast range of cellular processes including specification of epithelial cell identity, neural and cardiac development as well as stem cell differentiation (Mazin et al., 2021; Park et al., 2020; Wright et al., 2022). Alternative splicing is ubiquitous in mammalian cells – it is estimated that around 90% of all multiexon genes in humans have more than one transcript isoform (Wang et al., 2008). Further underscoring its importance, dysregulation of alternative splicing is common in diseases such as cancer and muscular dystrophies (López-Martínez et al., 2020).

Despite its importance, the role of alternative splicing in cell identity as well as a molecular indicator of cell type is not fully understood. Gene expression (as an aggregate of expression for all associated transcript isoforms) is well established as a robust indicator of cell identity (Dumitrascu et al., 2021; Pullin and McCarthy, 2024; Zeng, 2022). Groups of genes are often used as cell-type specific markers to delineate subpopulations and functional consequences of the altered expression of genes are relatively well annotated. In contrast, it is not certain if alternative splicing is of similar cellular importance and utility for cell classification. Historically, accurate quantification of transcript and/or splice event usage has been hampered by poor sequencing depth or probe coverage, as well as a lack of robust statistical or bioinformatic methods (Katz et al., 2010). Hence, most studies based on RNA sequencing or microarray data have derived estimations of aggregate gene expression without regard to the underlying splicing. However, with the advent of improved methods of detecting and quantifying alternative splicing, it is apparent that different transcript isoforms are associated with diverse functions in the cell (Marasco and Kornblihtt, 2022; Stamm et al., 2005). Indeed splice events may provide greater insights into cell identity than transcript expression, although this is yet to be formally demonstrated on a large scale.

The use of alternative splicing for data-driven methods such as cell type clustering and classification is emerging and has potential implications for clinical prognosis and disease subtype classification, notably in the context of cancer (Cai et al., 2020; Feng et al., 2021; Xiao et al., 2024). There has been evidence that supervised clustering methods using alternative splicing can potentially perform better than when using gene expression levels, but it is unclear as to why this may be the case, and whether alternative splicing data should be handled differently to gene expression (Johnson et al., 2018). There is a need for specialised bioinformatics tools for the processing of alternative splicing data and new methods in modelling splice inclusion levels and confounding variables in splicing data continue to be developed (Ascensão-Ferreira et al., 2024; Slaff et al., 2021). Despite the growing abundance of models tailored to alternative splicing, there is no concrete consensus as to what kind of model is the most suitable.

It is reported that splicing values are bimodal, meaning that cells often produce one or another of a given choice of alternative isoforms, rather than producing both simultaneously (Ascensão-Ferreira et al., 2024). This has been described from single-cell RNA-Seq experiments (Shalek et al., 2013) as well as bulk RNA-Seq experiments (Busch and Hertel, 2015; Tapial et al., 2017), and splice events exhibiting a clear bimodal distribution have been shown to be biologically relevant (Wan and Larson, 2018). However, it is debated whether bimodality is a true feature of the splicing landscape, or whether it simply arises from technical noise such as low sequencing coverage or intrinsic cellular behaviour unrelated to phenotype – such as transcriptional and spliceosomal kinetic variation (Najar et al., 2020; Wan and Larson, 2018).

To address the above questions, we analysed the RNA Atlas (Lorenzi et al., 2021), an ultra-deep RNA sequencing dataset obtained from 300 different types of cell and tissue samples across the human body. This allowed us to obtain a complete view of the human transcriptome. Through comparative gene expression and alternative splicing analysis, we were able to accurately compare the cell type distribution of splice events to that of gene expression. Our analysis shows that phenotypic differences in cells are faithfully reflected in their alternative splicing signatures and that alternative splicing in some cases performs better in data-driven cell type classifications. Further investigation into the distributions of splice event inclusion values and aggregate gene expression levels revealed that the zero-inflated and bimodal nature of splicing values is not due to sequencing or intrinsic cellular artifacts but is how splice events are inherently distributed amongst cell and tissue types. Our results provide deep insights into the way which transcriptomic information is expressed in splicing and how it fundamentally differs to aggregate gene expression.

## RESULTS

### A comprehensive map of human alternative splicing using the RNA Atlas

To obtain a comprehensive view of human alternative splicing, we analysed the RNA Atlas (Lorenzi et al., 2021), which contained ultra-deep short read sequencing data for 300 samples comprising of 45 tissue types, 166 primary cell types and 89 immortalised cell lines (Fig. 1a). Ribosomal RNA depletion (total RNA) libraries were used to prepare 294 samples and polyA capture libraries were used to prepare 295 samples (see Methods). Reads of 74bp in length were obtained at average sequencing depths of 135 million reads per sample for total RNA (59.7 billion reads altogether) and 58 million reads per sample for polyA capture (19.8 billion reads altogether).

**Figure 1.**
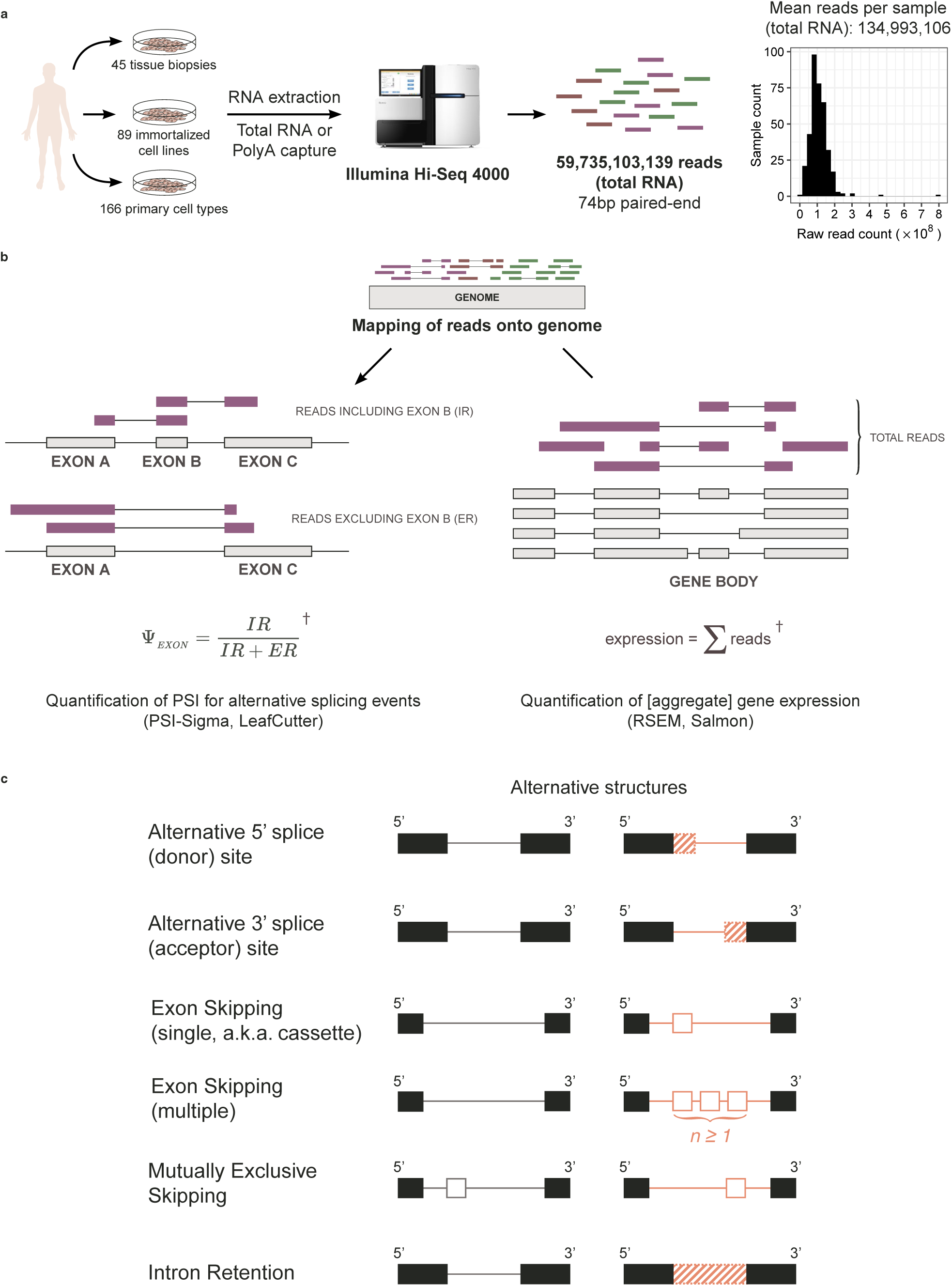
Comparative analysis of alternative splicing and aggregate gene expression using the RNA Atlas. **a)** Summary of the RNA sequencing process to generate the RNA Atlas. **b)** Illustration of bioinformatic workflow and disambiguation of alternative splicing and aggregate gene expression. ^†^Equations illustrate the basic principle of expression and splicing quantification and as such, the details of scaling, normalisation and distribution modelling commonly employed by bioinformatics algorithms are omitted. **c)** Summary of the common archetypal RNA structures generated by alternative splicing. Structures on left and right represent alternative exon connectivity for each mode of splicing. Black exons are constitutive.

To compare alternative splicing and aggregate gene expression (henceforth referred to as expression) in the RNA Atlas, we used two bioinformatics tools to align reads to the genome and two tools to analyse reads. Gene expression was quantified using RSEM or Salmon (in mapping-based mode) and alternative splicing was classified and quantified using PSI-Sigma or LeafCutter (Fig. 1b). These tools were chosen for each type of analysis because they employ different strategies, so any artifacts from tools should become evident in comparative analysis. Gene expression analysis yielded transcripts-per-million (TPM) values for genes annotated in Ensembl, and alternative splicing analysis yielded local Percent Spliced-In (PSI) values for events classified as one of six possible archetypes (PSI-Sigma) (Fig. 1c) or splice junctions (LeafCutter). For the total RNA dataset, there were 463,861 (PSI-Sigma), 194,073 (LeafCutter) splice events, and 50,026 (RSEM), 52,415 (Salmon) Ensembl genes detected in at least one sample. For the polyA capture dataset, there were 111,600 (PSI-Sigma), 158,620 (LeafCutter) splice events, and 44,242 (RSEM), 47,000 (Salmon) Ensembl genes detected in at least one sample.

### Alternative splicing strongly correlates with cell identity

For an overview of the expression and splicing landscapes, we generated UMAP projections based on the matrices of PSI and TPM values generated by each tool (Figs. 2a, d, g, h). Unsupervised clustering was performed by taking the consensus of a self-organising map ensemble, yielding 27 clusters (total RNA/PSI-Sigma), 37 clusters (total RNA/LeafCutter) and 36 clusters (total RNA/RSEM, totalRNA/Salmon), annotated as a colour overlay on the UMAP (see Methods). When clustering based on alternative splicing PSI values (PSI-Sigma (Fig. 2a), LeafCutter (Fig. 2d)), the proximity between similar cell and tissue types in the UMAP projection correlated well with cell type. Distinct clusters were well separated with median silhouette scores of 0.265 and-0.08 respectively – clear separation was observed between tissue biopsies, primary cells and cancer cell lines. Brain tissue (for the PSI-Sigma dataset, marked by an A5SS event in *PTBP2*, a known regulator of splicing (Fig. 2b) (Dawicki-McKenna et al., 2023)) and heart tissue samples were particularly distinctive and tightly grouped. In both datasets, cancer cell lines such as melanoma, T-ALL and neuroblastoma were assigned distinct clusters. In the PSI-Sigma dataset, the melanoma cluster was almost complete except for LOX IMVI cells. A single exon skipping event in *SMAD3* was a notable marker of the neuroblastoma cluster (Fig. 2c) (Boudreault et al., 2024). Other cell lines, including colon, bdreast and ovarian cancers were in mixed clusters. Similar primary cell types clustered together rather than according to organ of origin. Mesenchymal cells, fibroblasts and smooth muscle cells were broadly placed into two clusters. Strikingly, clusters of endothelial cells were complete for both PSI-Sigma and LeafCutter (the latter marked by an alternative last exon event in *FLT1* (Fig. 2e)), encompassing all available samples with no other cell types. The epithelial cell cluster in both PSI-Sigma and LeafCutter was almost complete except for amniotic and retinal epithelial cells, suggesting they could be distinct sub-types. Epithelial cells were distinguished by a multiple exon skipping event in Tight Junction Protein 1 (*TJP1*), which encodes for ZO1, a protein that stabilises tight junctions (Fig. 2f) (Fanning and Anderson, 2009). These results show that distinct cell and tissue types are distinguished by their alternative splicing patterns, with some (such as brain and heart tissue, endothelial and epithelial cells) being more distinct than others (such as mesenchymal cells, fibroblasts and smooth muscle cells).

**Figure 2.**
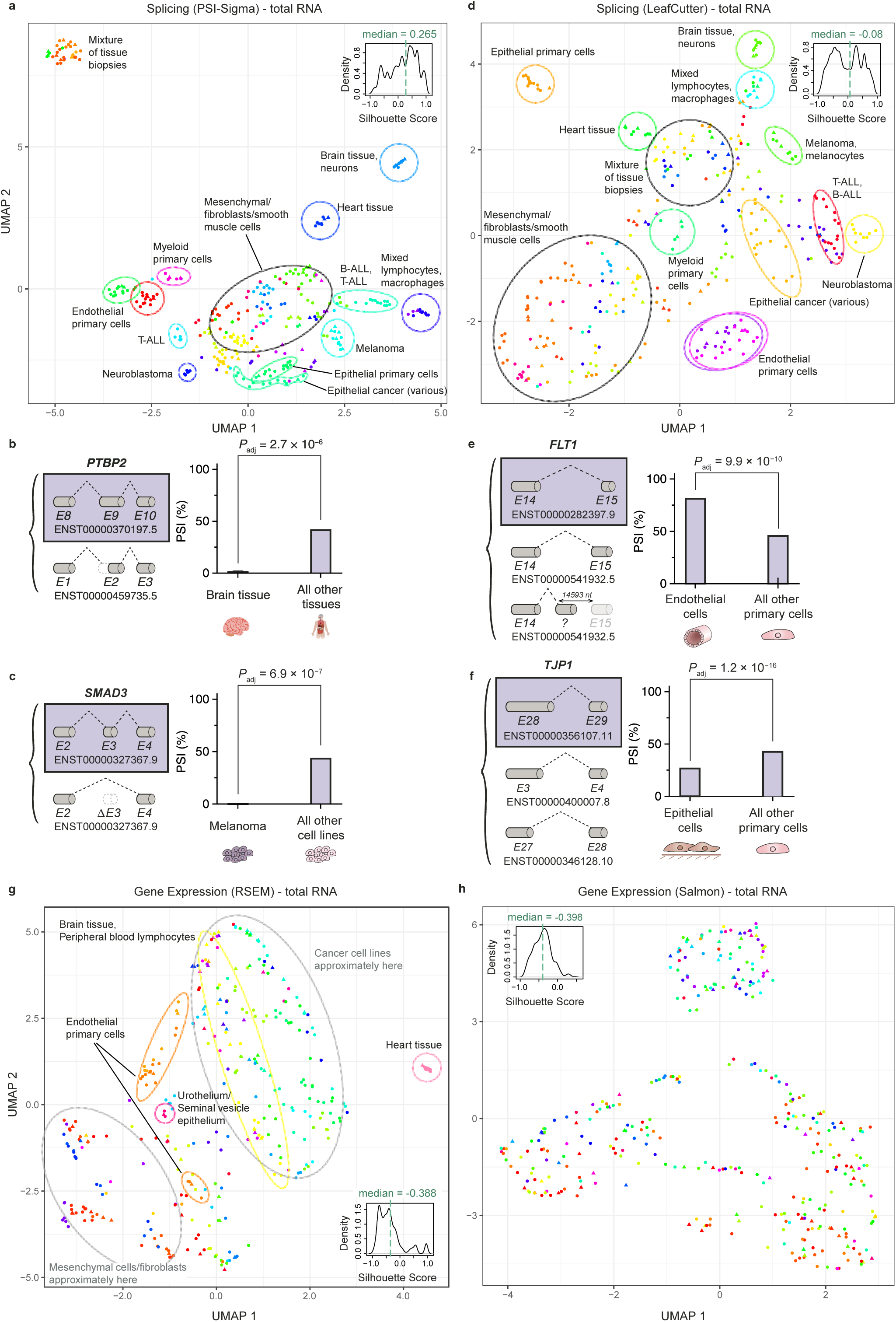
Splicing and gene expression, from the same samples, generate different unsupervised clusters. **a)** UMAP plot based on splicing PSI values detected by PSI-Sigma for the total RNA dataset. Each data point represents one RNA-Seq sample. Silhouette score distributions shown in insets - higher scores reflect greater cluster separation. **b-c)** Notable markers of splicing for select UMAP clusters shown in a). **b)** Splicing marker of brain tissue, an alternative splice acceptor (A3SS) event in PTBP2, matching to transcript ENST00000370197.5. **c)** Splicing marker of multiple melanoma subtypes, a multiple exon skipping event in SMAD3, matching to transcript ENST00000327367.9. **d)** UMAP plot based on splicing PSI values detected by PSI-Sigma for the total RNA dataset. Each data point represents one RNA-Seq sample. Silhouette score distributions shown in insets - higher scores reflect greater cluster separation. **e-f)** Notable markers of splicing for select UMAP clusters shown in d). **e)** Splicing marker of various epithelial cells throughout the body, an alternative 5’ splicing event in TJP1, matching to transcript ENST00000356107.11. **f)** Splicing marker of various endothelial cells throughout the body, an alternative last exon event in FLT1, matching to transcript ENST00000282397.9. The bottom splice junction maps to an unannotated exon in between exon 14 and 15 of ENST00000541932.5. **g-h)** UMAP plots based on gene expression values detected by RSEM or Salmon for the total RNA dataset. Each data point represents one RNA- Seq sample. Silhouette score distributions shown in insets - higher scores reflect greater cluster separation.

Clustering and embedding by gene expression TPM values (RSEM (Fig. 2g) and Salmon (Fig. 2h)) did not group most samples in a clear manner, with clusters displaying a high degree of overlap. For RSEM, heart tissue samples were embedded separately from the rest of the samples, but unrelated cell types such as brain tissue, melanoma and colon cancer were in proximity. All clusters arising from Salmon-based read alignments were found to overlap, hence we omitted annotations from the figure. Indeed, the median silhouette scores were-0.388 and-0.398, with all Salmon-based clusters notably having scores lower than 0.5. Given that we found that some samples were clustered correctly for the Salmon dataset despite their unclear spatial embedding, this is not an error with Salmon quantification but is indicative of reduced UMAP embedding power. The differences in clustering and embedding power between splicing and expression were recapitulated in the polyA capture dataset. Both PSI- Sigma and LeafCutter splicing data (median silhouette 0.225 and 0.044) (Supplementary Figures 1a, b) outperformed RSEM and Salmon expression data (median silhouette 0.032,-0.465) (Supplementary Figures 1c, d). Additionally, the median silhouette scores associated with tSNE plots for all total RNA and polyA capture data were consistent with those for UMAP plots, with improved embedding performance with splicing data compared to expression data (Supplementary Figure 1e, tSNE plots not shown). These data suggest that alternative splicing data confers more robust embedding using dimensionality reduction tools than expression levels.

### Alternatively spliced genes are associated with distinct functional outcomes

We sought to understand the differences in splicing between each UMAP cluster and whether differentially spliced genes are indicative of cell phenotype. For each UMAP cluster, differential splicing and expression analyses were performed based on one-vs-rest comparisons of the same sample type (i.e. cell line, primary cell or tissue biopsy) (see Methods). Overall, there were 12,717 (PSI-Sigma), 10,192 (LeafCutter) differential splicing events detected from total RNA, and 2,324 (PSI-Sigma), 7,302 (LeafCutter) events from polyA-captured RNA (Supplementary Figure 2a).

Evaluating gene expression alone, there were 39,900 (RSEM), 25,689 (Salmon) genes differentially expressed in total RNA and 36,448 (RSEM), 14,458 (Salmon) from polyA-captured RNA (Supplementary Figure 2a). Despite LeafCutter being unable to detect intron retention events, the concordance between LeafCutter and PSI-Sigma differentially spliced genes remained high (7,410 genes). In both PSI-Sigma total RNA and PSI-Sigma polyA capture datasets, intron retention (IR) events were the most common (66.8% for total RNA, 47.2% for polyA capture), followed by single exon skipping (SES) (13.3% for total RNA, 3.6% for polyA capture) (Supplementary Figures 2b, c). Splice events occurred most frequently within 3’ UTRs (37.0% for total RNA, 19.7% for polyA capture) and protein-coding regions (38.0% for total RNA, 18.4% for polyA capture). While intron retention events usually occurred within protein-coding regions and 3’ UTRs, SES and MES events almost always occurred in protein-coding regions. These patterns remained consistent across cell and tissue types.

Differentially expressed and differentially spliced genes were analysed in the context of Gene Ontology. Clusters (as described in Figs. 2a, d, g, h) were often associated with cell-type specific GO terms such as “G2/M transition of mitotic cell cycle” for neuroblastoma cell lines, “growth cone” for brain tissue/neurons, “angiogenesis” for endothelial primary cells and “cell-cell junction” for epithelial primary cells (Supplementary Figure 2d, e). However, there were also more general cellular processes common to multiple clusters such as “actin cytoskeleton”, “alternative mRNA splicing, via spliceosome”, “Golgi apparatus” and “protein serine/threonine kinase activity”. This prompted us to explore whether differentially spliced genes were only associated with a subset of GO terms. Fig. 3 shows a network of the Gene Ontology in its entirety, with nodes colour coded according to the statistical significance of their enrichment in total RNA/PSI-Sigma genes compared to total RNA/RSEM genes. Differentially spliced genes were most often enriched for GO Terms also associated with differentially expressed genes, generating purple nodes. Many of these were large nodes (associated with a greater number of genes), such as “endoplasmic reticulum” (CC), “transcription factor complex” (CC), “RNA binding” (BP), “protein binding” (BP), “protein ubiquitination” (BP) and “protein transport” (MF). These GO terms were associated with a high number of genes in the database and so are general in their description of biology. Smaller GO terms associated with fewer genes and enriched in the PSI-Sigma differential splicing dataset (generating red nodes) were concentrated around certain hubs in the GO term network. For example, the neighbourhood of “hydrolase activity” (MF) and “transcription factor complex” (CC) nodes had clusters of other small GO terms enriched in PSI-Sigma genes. Nodes in the GO term network are closer to each other if their GO terms describe similar sets of genes, thus the data suggests that splicing alone regulates a limited subset of cellular functions. In stark contrast, GO terms enriched for RSEM differentially expressed genes were more common and widespread regardless of the size of the GO terms since almost all small nodes were coloured cyan. Some GO terms were solely associated with differentially expressed genes, such as those in the neighbourhood of “anatomical structure development” (BP), “regulation of gene expression” (BP) and “developmental process” (BP). Furthermore, distribution of GO terms enriched in differentially spliced or expressed genes remained consistent for all comparisons between PSI-Sigma, LeafCutter, RSEM and Salmon for both the total RNA and polyA capture dataset (Supplementary Figure 3). Given that there were a higher proportion of sample-specific GO terms in the splicing data (63% of GO terms *P* < 0.01 enriched in only one cluster in total RNA/PSI-Sigma compared to 44% for total RNA/RSEM), it could be that the functional consequences of splicing are more selective. It is possible that alternative splicing does not inherently modulate a wide range of biological pathways despite a higher proportion of differentially spliced genes (89% of total RNA/PSI-Sigma genes) than differentially expressed genes (44% of total RNA/RSEM genes) showing significant enrichment for at least one GO term in any cluster. Indeed, it has been proposed that alternative splicing could be non-functional (Pickrell et al., 2010; Reixachs-Solé and Eyras, 2022). Equally, but more difficult to quantify, the differences in enrichment between splicing and expression here may also reflect insufficient annotation of splicing consequences in the GO database.

**Figure 3.**
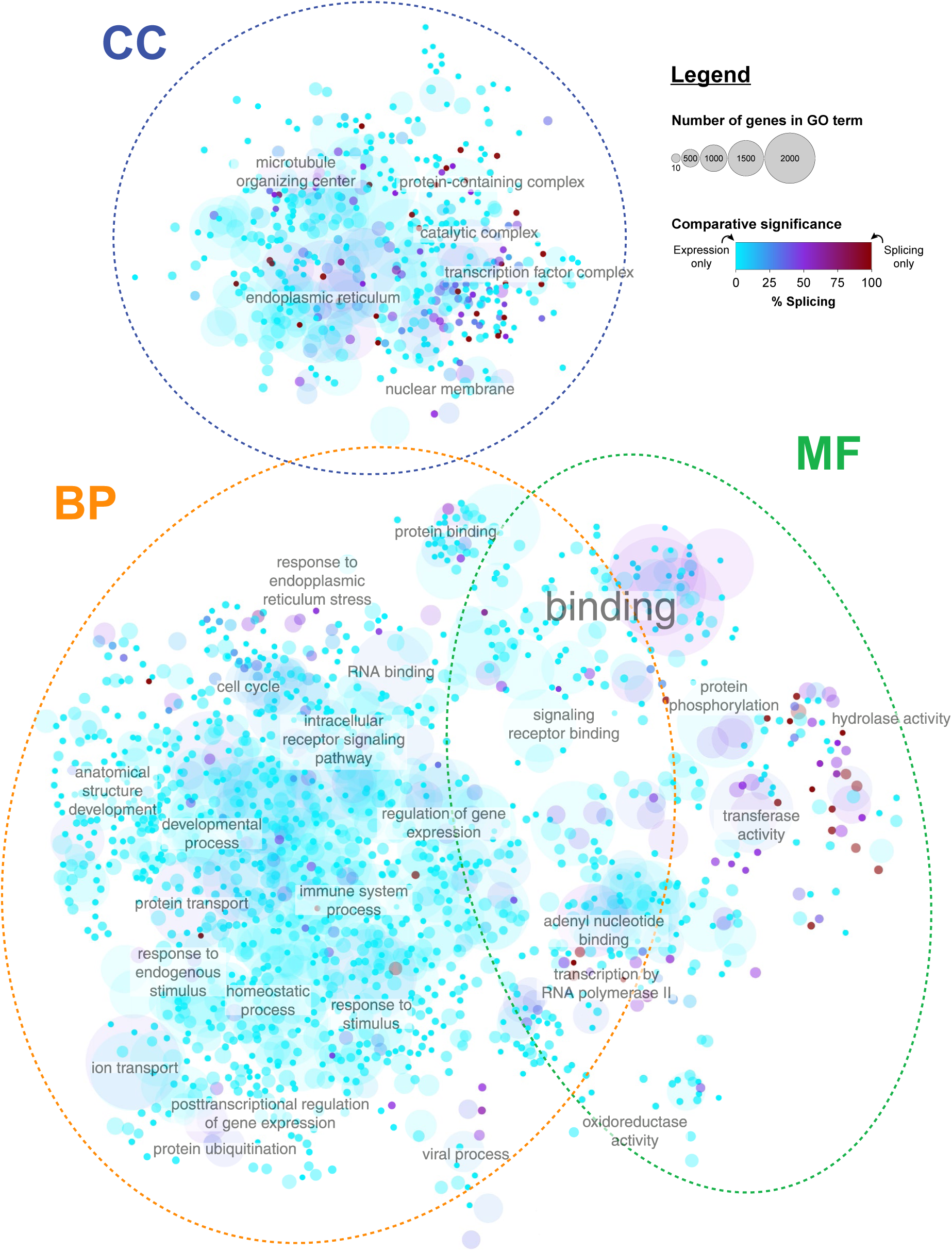
Alternative splicing is associated with a restricted repertoire of functions. The complete Gene Ontology network showing nodes enriched for differentially spliced genes (total RNA, PSI-Sigma) and differentially expressed genes (total RNA, RSEM). Each node represents a GO term. The size of each node reflects the number of genes associated with the GO term – GO terms with more genes have a larger node. Only the names of select nodes of interest are shown. Color of each node indicates the percentage of the combined – log_10_ FDR attributed to the PSI-Sigma dataset. Blue colors: PSI-Sigma significance lower than RSEM; purple: equal statistical significance; red colors: PSI-Sigma significance greater than RSEM. BP: Biological Process; CC: Cellular Component; MF: Molecular Function.

### Alternatively spliced non-coding RNAs contribute to better cell classification

Given clear differences between the alternative splicing and gene expression landscapes of cells and that these differences are functionally significant, we explored if alternative splicing data yields different results than expression data for the training and supervised classification of cell and tissue types. To perform unsupervised clustering, we utilised four models in the Python scikit-learn toolkit: C-Support Vector Classification (svm), k-Nearest Neighbors Classifier (knn), Logistic Regression (logreg), and Decision Tree Classifier (decisiontree). Alternative splicing PSI (PSI-Sigma, LeafCutter) or gene expression TPM (RSEM, Salmon) values from each biological sample of the RNA Atlas were labelled according to the unsupervised clustering assignments that we generated in Figs. 2a, d, g, h and Supplementary Figures 1a-d. Model accuracy and precision was evaluated by randomly sampling (bootstrapping) a varied percentage of samples to be withheld for validation. Percentages ranged from 5% to 90% with the remainder used for model training, and 1000 bootstraps were done at every 5% increment. This was repeated separately for primary cells, tissue biopsies and cell lines.

Receiver-Operator Characteristic (ROC) curves were generated for each classification (not shown). Since larger training sets are preferred for the practical identification of unknown samples, we elucidated ROCs of the classification models at a percentage withheld of 10%. Sample labels related to multiple UMAP clusters, hence the task of classification was treated as a multi-class problem. ROC curves and their associated mean Area Under the Receiver-Operator Characteristics (AUROCs) (Hand and Till, 2001) were generated for each model and dataset, shown in Fig. 4. Overall, classification was robust across all cell and tissue datasets for svm, logreg and knn, with small differences in AUROC between datasets and classification models. Decisiontree was consistently the poorest performing classifier. For the total RNA dataset, alternative splicing almost always outperformed gene expression for classification of cell lines, primary cells and tissue biopsies, with the exception being cell lines using knn. Splicing total RNA data attained the six highest average AUROC values in the analysis: cell line/logreg (0.918), primary cells/svm (0.906), cell line/svm (0.904), tissue biopsy/svm (0.899), primary cells/logreg (0.895) and tissue biopsy/logreg (0.889). In contrast, less consistent results were observed in the polyA dataset, with expression data outperforming splicing data in 6 of 12 analyses – here, classifications based on splicing and expression achieved the two highest AUROC values: splicing/cell line/logreg (0.871) and expression/cell line/svm (0.867). Whilst splicing fared better in all tissue biopsy classifications for total RNA, expression fared better for all tissue biopsy classifications for polyA capture.

**Figure 4.**
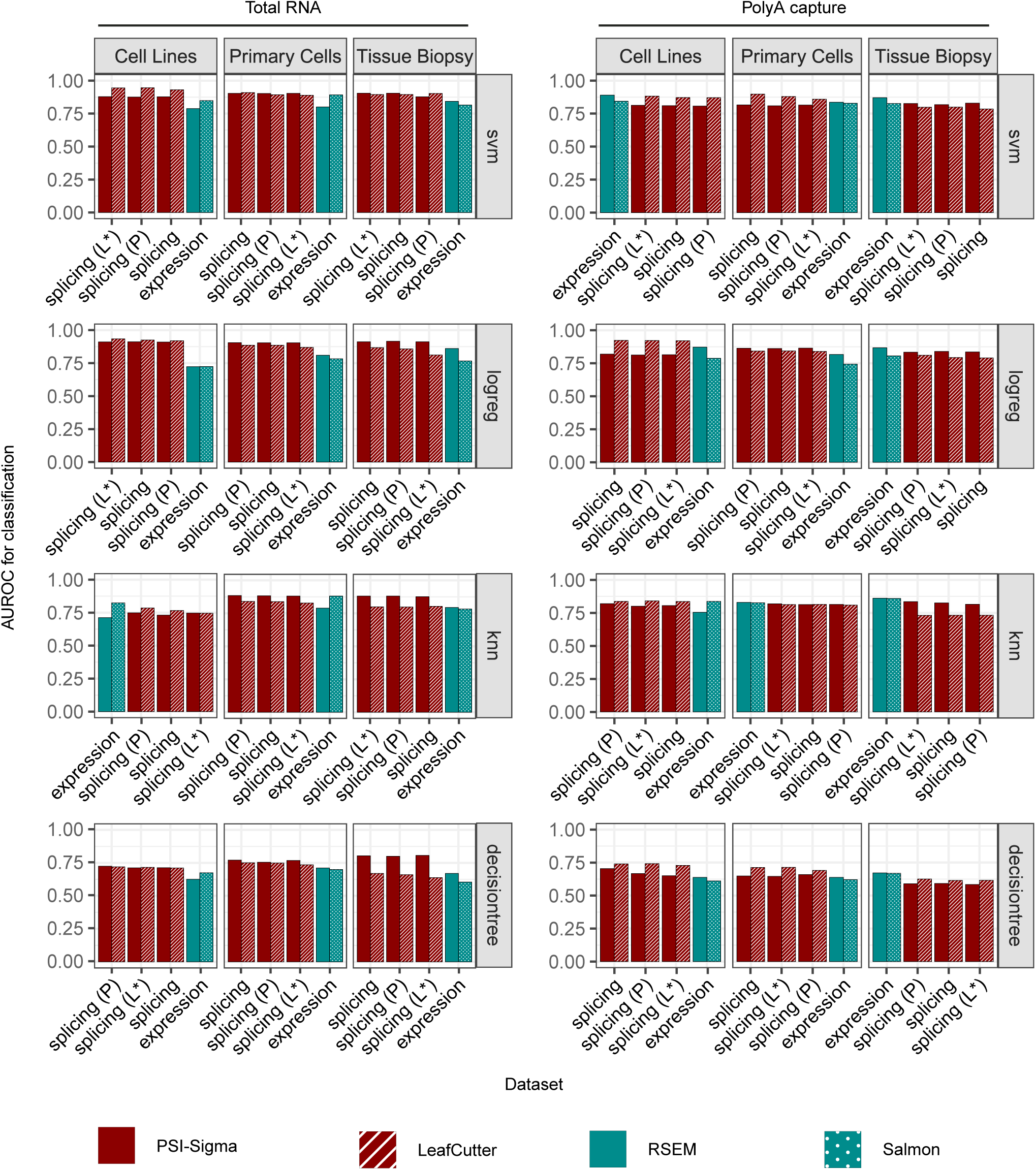
Non-coding transcripts are required for improved classification power of cell lines, primary cells and tissue biopsies using alternative splicing. Area Under the Receiver-Operator Characteristics (AUROCs) for the total RNA and polyA capture datasets. Pairs of bars representing the same dataset (i.e. splicing or expression) are sorted in decreasing order of their average from left to right. Bar plots are coloured according to the mode of analysis used for classification: solid dark red (PSI-Sigma), striped dark red (LeafCutter), solid teal (RSEM), teal with white dots (Salmon). L*: splice events which do not match to Ensembl lncRNA transcripts only, P: splice events matched to protein-coding transcripts only, svm: C-Support Vector Classification, logreg: Logistic Regression, knn: k-Nearest Neighbours Classifier, decisiontree: Decision Tree Classifier.

To explore whether splicing data yields better classification accuracy when there are less data available for training, we sampled classification accuracy at multiple percentage withheld values (percentage of dataset not used for training) with the results shown in Supplementary Figures 4a, b. The accuracy of classification by splicing was much higher than that for expression, both in terms of AUC values (increase of 0.189 for total RNA, 0.087 for polyA capture) as well as maximal attainable accuracies at percentage withheld values less than 20%. This and the above results suggest that sample classification via transcriptomic data can be improved by using alternative splicing data instead of expression data, especially using total RNA libraries instead of polyA capture libraries.

We explored the possibility that non-coding RNAs could account for the differences in classification power described above. Since the total RNA dataset was prepared via ribosomal RNA depletion, reads could have originated from polyadenylated or non-polyadenylated non-coding RNA molecules (Lorenzi et al., 2021). In particular, long non-coding RNAs (lncRNAs) were of interest because they are relatively well annotated in Ensembl. To that end, we explored whether the removal of lncRNAs from the total RNA data could affect the classification power of alternative splicing data. The classification bootstrapping process was repeated on the data with lncRNA-associated splice events removed (L*) as well as that with only protein-coding associated splice events (P) (see Methods). The number of non-lncRNA splice events were 458,052 (total RNA, PSI-Sigma), 110,335 exons (polyA capture, PSI-Sigma) and 55,783 (total RNA, LeafCutter), 54,784 junctions (polyA capture, LeafCutter). The number of protein-coding only splice events was 449,089 (total RNA, PSI-Sigma), 106,091 exons (polyA capture, PSI-Sigma) and 100,419 (total RNA, LeafCutter), 98,963 junctions (polyA capture, LeafCutter).

The AUROCs for L* (lncRNA-associated splice events removed) and P (only protein-coding associated splice events) datasets are shown in Fig. 4. If lncRNAs or other non-coding RNAs are responsible for differences in classification power between splicing and expression, then the AUROCs from L* and P data should take on intermediate values between that of unmodified splicing and expression data. Of the 12 classifications in total RNA data, 4 saw intermediate performance of L* and P datasets, 5 saw mixed performance where one fared better than the unmodified splicing dataset and the other fared worse, and 3 saw L* and P datasets perform even better than the unmodified splicing dataset which had already outperformed the expression dataset (however, these differences are small). In the polyA capture data, 6 classifications saw intermediate performance of L* and P datasets, 5 classifications with mixed performance and 1 classification where L* and P performance was less similar to expression than the unmodified splicing dataset (cell lines/knn). It is likely that removing splice events associated with non-coding RNA reduces the accuracy and precision of classification due to there being less information available for training. Indeed, for 4 of 6 classifications across both total RNA and polyA capture datasets where unmodified splicing data was outperformed by gene expression, the L* and P datasets displayed mixed performance, whereas the remaining 2 classifications displayed intermediate performance of L* and P datasets, with small improvements upon unmodified splicing data. These results do not suggest that non protein-coding RNAs present in splicing data represent an incremental improvement upon the classification power achievable using expression data. Rather it points to splicing containing an independent set of information with a distinct underlying structure.

### Cell-type specific features define the landscape of alternative splicing

We suspected that the differences in splicing and expression landscapes thus far could be due to differing distributions of PSI or TPM values, including the distribution of zero values. To quantify cell usage of alternative splicing and gene expression, we devised a metric which we call “feature degeneracy”. Given a dataset containing *m* features (i.e. splice events or genes), values are recorded across *n* samples (Fig. 5a). For the RNA Atlas, *n* is 294 (total RNA) or 295 (polyA capture). As a simple way to measure the presence or absence of a feature in a sample, each zero value of PSI or TPM is set to 0 and each non-zero value is set to 1. For a given feature, we define a *degeneracy factor* (*d*) which corresponds to the number of non-zero values. When *d* > 1, the feature is considered “degenerate” – it is re-used as a means of control in multiple cells and is hence unspecific to any single cell or tissue type, irrespective of any functional significance. When *d* = 1, there are no degenerate values and when *d* = *n*, all values are degenerate (Fig. 5b).

**Figure 5.**
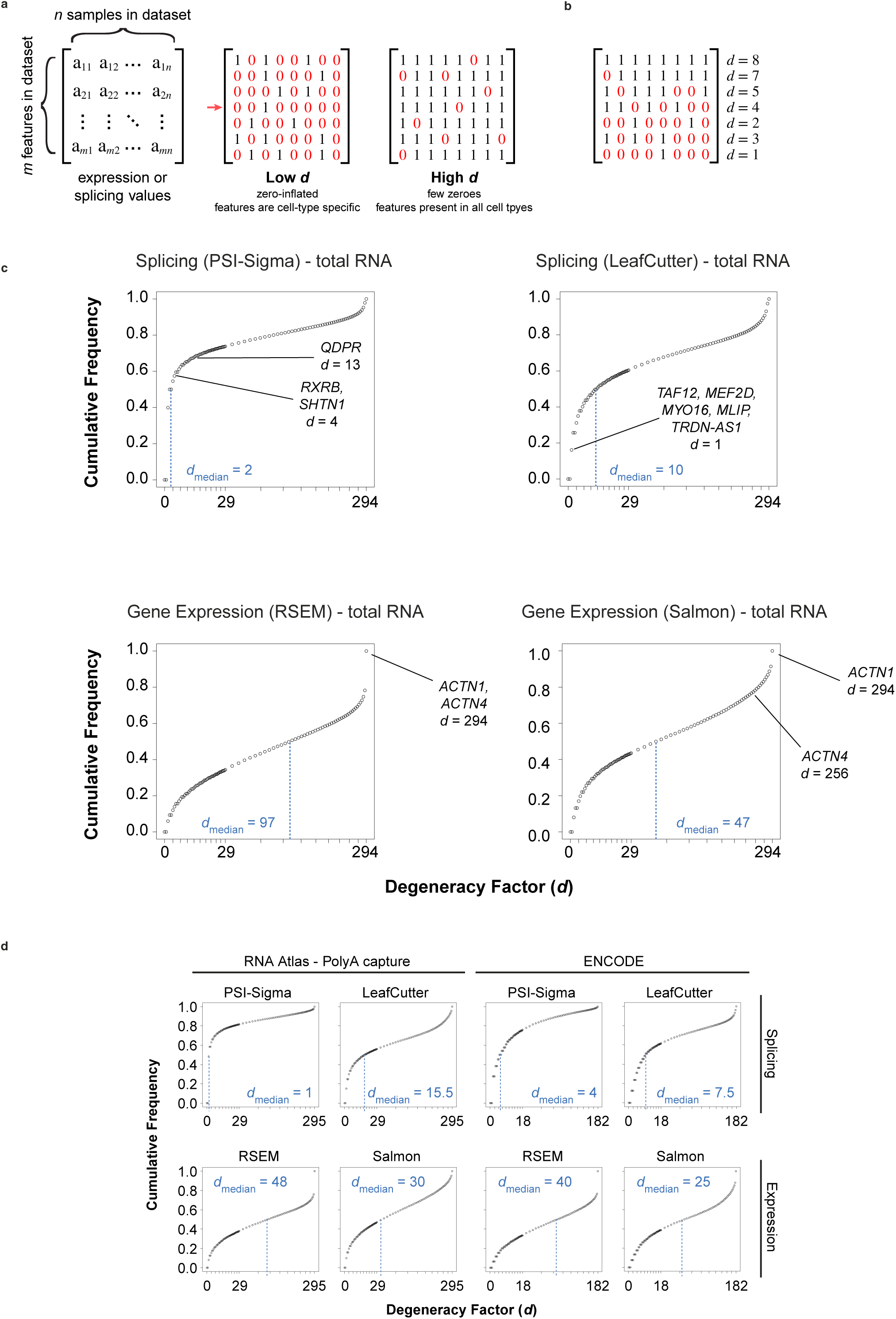
Degenerate features distinguish gene expression from alternative splicing. **a)** A degeneracy factor (*d*) is measured individually for each feature. The row indicated by the red arrow signifies a feature with no degeneracy. **b)** Example *d* values for example features. **c, d)** Cumulative frequency plots of Feature Degeneracy Factors (*d*). Features were measured from the alternative splicing PSI levels (PSI-Sigma, LeafCutter) or the gene expression TPM values (RSEM, Salmon) for total RNA or polyA capture datasets of the RNA Atlas in addition to the complete ENCODE RNA- Seq dataset. Median *d* values of each dataset are indicated below the vertical dotted blue lines. **c)** RNA Atlas total RNA datasets. Genes of note with high or low *d* values are annotated. \**d* value is relatively high because the splice event in *QDPR* was present exclusively in 9 of the 12 total distinct melanoma samples. **d)** RNA Atlas poly A capture and ENCODE RNA-Seq datasets.

Degeneracy factors were calculated for each feature in the RNA Atlas, and plotted as cumulative frequency distributions in Figs. 5c, d. In the total RNA dataset, splicing features displayed low degeneracy, with median *d* values of 2 (PSI-Sigma) and 10 (LeafCutter). In contrast, gene expression values had higher median *d* values of 97 (RSEM) and 47 (Salmon) (Fig. 5c). For the polyA capture dataset, differences between splicing and expression were consistent, with median *d* values of 1 (PSI-Sigma), 15.5 (LeafCutter), 48 (RSEM) and 30 (Salmon) (Fig. 5d). This data shows that splice events are more likely to only be found in a subset of cell types. Indeed, splice events with low *d* values are of biological importance: in the total RNA/PSI-Sigma dataset, *QDPR*, known to be associated with melanoma, had a splice event found exclusively in 9 of 12 available melanoma samples; *RXRB*, a crucial nuclear receptor found in the brain and *SHTN1*, a key regulator of axonogenesis, were each found solely in brain and neuron tissue samples (Fig. 5c). In the total RNA/LeafCutter dataset, multiple regulators of cardiomyocyte differentiation *TAF12, MEF2D, MYO16, MLIP* and *TRDN-AS1* were found exclusively in heart samples (Fig. 5c). In comparison, the higher overall degeneracy of gene expression shows that most genes are expressed in all cell types and tissues. Indeed, housekeeping actinin genes *ACTN1* and *ACTN4* had degeneracy factors of (294 and 294 respectively, total RNA/RSEM and 294 and 256 respectively, total RNA/Salmon) (Fig. 5c). To corroborate these observations on an independent dataset of comparable scale, we also calculated the degeneracy factors from the ENCODE Project (Luo et al., 2020), totalling 182 samples with a read count per sample greater than 100 million paired-end reads (Fig. 5d, see Methods). This yielded median *d* values of 4 (PSI-Sigma) and 7.5 (LeafCutter) for splicing, and values of 40 (RSEM) and 25 (Salmon) for expression, confirming observations with the RNA Atlas. Thus, the presence or absence of a splice event is more likely to be a unique marker of cell type compared to the presence or absence of genes themselves.

We further explored the quantitative differences in the distributions of PSI and gene expression levels. Features can be thought of as control dials with a fixed set of increments defined by the cell (Fig. 6a). PSI and expression values obtained from RNA-Seq approximate these increments with variance due to sequencing artifacts, random noise and tolerances in the intracellular regulatory network. We asked the question of whether increments occur only at a few positions (fewer increments, greater increment distances) (Fig. 6b) or at a diverse range of positions along the range of observed feature values (more increments, smaller increment distances) (Fig. 6c). We made the assumption that larger changes in the level of a feature is likely to have a stronger impact on cell identity than a smaller change. Following this logic, if observed feature values are sufficiently similar amongst a set of samples, then the feature values are also degenerate in the sense that cells are employing the same strategy to define distinct molecular phenotypes.

**Figure 6.**
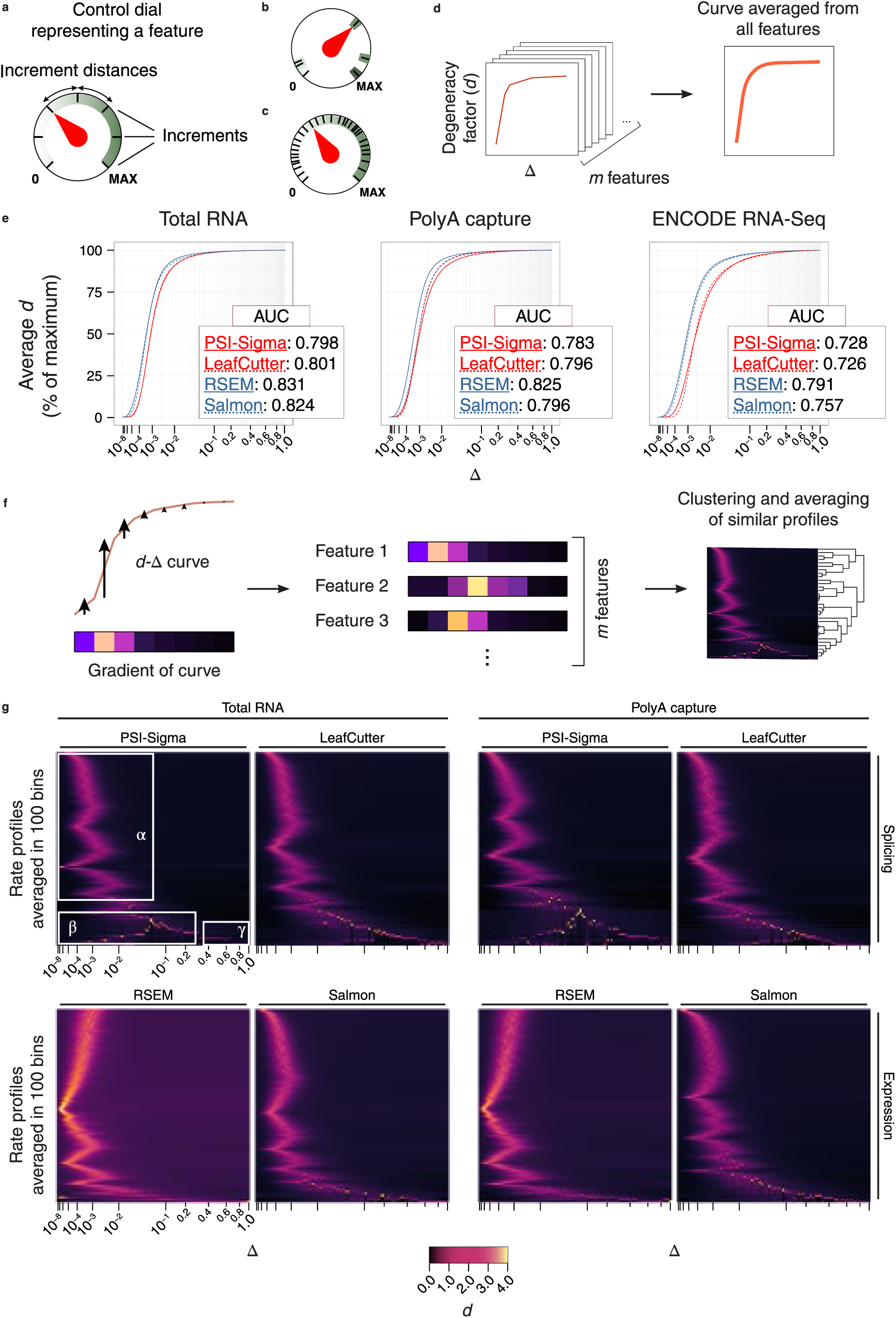
Gene expression levels display finer variation than alternative splicing. **a)** Features such as gene expression and splicing levels as control dials. **b)** A feature with fewer increments. **c)** A feature with more increments. **d)** Process of generating averaged *d*-Δ curves. **e)** Averaged *d*-Δ curves for each dataset. Red solid line: PSI-Sigma, red dotted line: LeafCutter, blue solid line: RSEM, blue dotted line: Salmon. **f)** Process of generating heatmaps based on the gradient of the *d*-Δ curves. **g)** Heatmap of gradient profiles of *d*-Δ curves. There are 100 rows each representing a bin of averaged feature profiles. Each column represents a single Δ value.

We employed a simple approach to survey the density of feature values by testing many possible noise tolerances across the full range of each feature. To accommodate for the more general definition of feature degeneracy, we expanded the definition of *d* (Supplementary Figure 5). The maximum value of each feature was scaled to 1 and values are converted to an interval with range ±Δ (the noise tolerance), such that intervals may overlap if their feature values are close together. When Δ = 0, the number of intervals is equivalent to the number of unique non-zero feature values (*n_zero_*) and the degeneracy factor is at a minimum value. As the value of Δ increases, more intervals merge, and the degeneracy factor increases since overlapping intervals are considered degenerate. Eventually, as Δ → 1, all intervals merge into one and degeneracy factor reaches a maximum. Thus, the new definition of *d* is the number of interval merges at a given value of Δ:

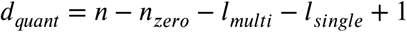

Where *l_multi_* is the number of intervals merged from multiple feature values, and *l_single_* is the number of unmerged intervals.

For each feature in the RNA Atlas and ENCODE datasets (Luo et al., 2020), quantified by the various splicing and expression analysis tools, a degeneracy-delta (*d*-Δ) curve was calculated for 96 values of Δ and averaged (as per Fig. 6d, see Methods). Fig. 6e shows the resulting curves. Degeneracy factors displayed increases starting from a Δ value as low as 1.0 × 10^−7^, indicating that a large proportion of non-zero values were close together. RSEM and Salmon datasets displayed sharp increases in degeneracy as Δ increased further, reaching more than 90% of their maximum *d* values at Δ values around 4.0 × 10^−3^ (total RNA and polyA capture). This means that the expression values of most genes are separated by less than 0.4% of their range across cell types. In comparison, PSI-Sigma/LeafCutter datasets did not reach 90% of their maximum *d* values until Δ values reached approximately 0.01. RSEM and Salmon consistently achieved higher AUC values than PSI-Sigma and LeafCutter, which indicates a higher overall degeneracy for expression values. In fact, degeneracy values were equal to or higher in RSEM and Salmon for all values of Δ, hence gene expression levels are closer together (denser) than splicing levels.

Having measured the density of feature values, we sought to understand the nature of their increment distances. To do this, we used the gradient of the *d*-Δ curve, which is proportional to the number of increment distances that are equal to the difference between consecutive Δ values (Fig. 6f). Heatmaps in Fig. 6g each show 100 bins that summarise the different types of gradient profiles found in each dataset, where one row represents one bin. For each bin, the profile is an average from of multiple similar features as determined by clustering. Most bins were characterised by a smooth unimodal distribution (Fig. 6g, area corresponding to α in the left side of plots). This suggests that feature values are often approximately separated by a small mean distance less than 0.01, with random variance from the mean separation likely due to noise. On average, the mean separation for splicing occurs further to the right of the plot then gene expression, suggesting that the features in splicing usually have a wider minimum separation. For the PSI-Sigma and LeafCutter curves, around 10% of bins (Fig. 6g, area corresponding to β in bottom of plots) displayed one or more well-defined peaks. These represent feature values which are present in few cell types and have one or more modal separation distances.

Some bins (Fig. 6g, area corresponding to γ in bottom right of plots) displayed peaks very close to Δ = 1, which means that features were either close to or equal to 0 or 1, with nothing in-between. These were more common in the splicing datasets as bins with peaks above 90% constituted 44% (total RNA/PSI-Sigma), 26% (total RNA/LeafCutter), 13% (total RNA/RSEM), 13% (total RNA/Salmon), 55% (polyA capture/PSI-Sigma), 20% (polyA capture/LeafCutter), 18% (polyA capture/RSEM) and 17% (polyA capture/Salmon) of all features. Together, these results show that the distribution of features for splicing PSI values is fundamentally different to that of expression TPM values in that the minimum separation between PSI values is wider and has a higher tendency to be either completely present in some cell types and completely absent in others.

The difference in feature value distribution between splicing and expression has fundamental implications on the way they relate to cell identity. For alternative splicing, this means that cell and tissue types tend to be distinguished by the sheer presence or absence of certain splice events. PSI levels of splice events are also less continuous and contain more outliers – akin to a control dial with few intermediate settings (Fig. 7). For gene expression, the mRNA of most genes remains present in most cell and tissue types but are regulated at different levels. Gene expression levels tend to be more fine-tuned, taking on a diverse range of values, akin to a control dial with many increments (Fig. 7). These differences provide a possible explanation as to why splicing and expression perform differently in data-driven clustering, classification and functional analyses, and may inform future treatment of transcriptomic data that is tailored to the underlying data structure.

**Figure 7.**
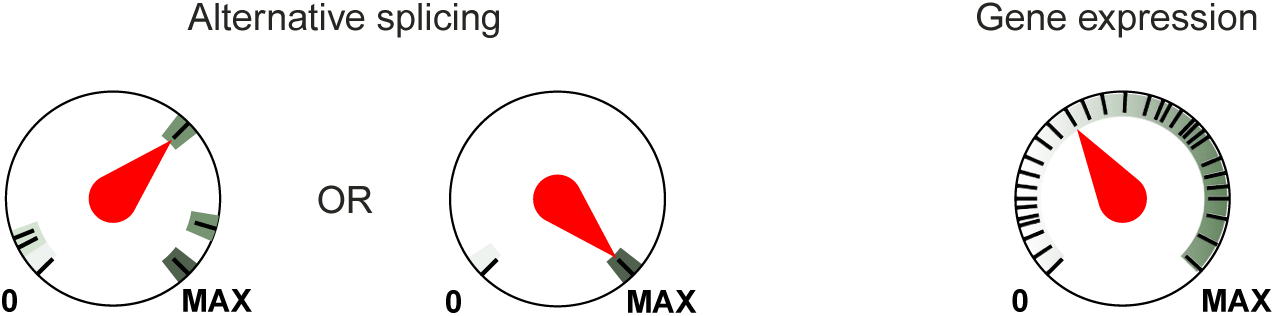
**Alternative splicing and gene expression show fundamentally distinct feature distributions.**

## DISCUSSION

Insights into the landscape of human alternative splicing are expanding as existing sequencing technologies mature and new technologies emerge. With the sheer scale and breadth of the RNA Atlas (Lorenzi et al., 2021), both in terms of the number of cell and tissue types in the dataset, as well as the sequencing depth and having both total RNA and polyA capture, we were able to construct a comprehensive picture of human alternative splicing. The ENCODE project (Luo et al., 2020), roughly twice in size and largely complementary in terms of cell and tissue types, served as a crucial dataset for independent validation of the insights obtained from the RNA Atlas.

A longstanding question has been whether alternative splicing has a significant role in determining cell identity. Our results suggest this to be true. By using splice inclusion values instead of gene expression values, we saw striking improvements in performance of UMAP embedding, unsupervised clustering and supervised classification of cell types, tissue types and cell lines. Importantly, we used orthogonal bioinformatics pipelines to quantify splicing (PSI-Sigma, LeafCutter) and gene expression levels (RSEM, Salmon) but our results remained consistent. These results align line with those from a previous large-scale study which showed that transcript-level expression data (as measured by Cufflinks v2) for mice and human Cancer Genome Atlas RNA-Seq samples generally outperformed expression level data for supervised classification (Johnson et al., 2018). For this study, we additionally found the improvement in classification performance when using splicing data was most evident for total RNA samples than polyA capture samples. Although the simplest hypothesis is that the presence or absence of non-polyadenylated RNAs (which are mostly non-protein coding) is responsible for this difference in classification power, this was not supported by our data. We propose an alternative explanation that the differences in classification power can be explained by splice events having lower degeneracy than the expression of genes and furthermore, the difference in degeneracy is more pronounced with data for total RNA. Low degeneracy means that the presence or absence of a splice event can be a clearer indicator of distinct cell types than small adjustments in the expression of multiple genes. As a result, distinct cell types would be more readily discernible by their splicing patterns than their gene expression patterns. Our data supports previous reports in literature that alternative splicing constitutes a layer of cellular information independent to gene expression, as well as recommendations to include alternative splicing analysis wherever possible in RNA-Seq studies, at least as a complementary analysis to gene expression, to gain deeper biological insights (Dam et al., 2023; Ha et al., 2021). Further studies are needed to determine if total RNA-derived alternative splicing data can yield superior classification power in a clinical context, especially if more specialised and comprehensive training datasets are used.

In general, splice events have a low degeneracy factor, with wide gradations in PSI levels. A subset of splice events possess a bimodal distribution of PSI levels. In previous studies in literature (mostly single-cell RNA-Seq) that demonstrated the presence of bimodality in the splicing landscape, the sequencing coverage has been the subject of scrutiny, with simulations showing that sampling bias can give the appearance of bimodality (Najar et al., 2020). However, the sequencing in the RNA Atlas is extremely deep, with an average of 135 million reads per sample for total RNA and 58 million reads per sample for polyA capture. Due to the RNA Atlas being a bulk sequencing dataset, it is also robust against cell-to-cell variations in transcriptional and co-transcriptional kinetics such as bursting and delayed splicing. Although degeneracy can be deflated for splice events with low expression levels, a very conservative minimum read count filter was applied before PSI-Sigma/LeafCutter (PSI values with numerator > 15 only), and RSEM/Salmon analyses (TPM values > 0.5 only) (see Methods), which minimises the impact of read noise on splice events and genes that have low expression.

Accounting for all these factors, the persistence of bimodality in our splicing data indicates that it is an inherent feature of the splicing landscape. Future studies involving quantitative analysis of alternative splicing should take into account or leverage the low degeneracy of splicing values.

We showed that splicing correlates well with cell type differences, however the functional significance of cell type-specific splicing patterns were not explored in this study as such an endeavour would involve analysis of upstream regulation of splicing or of downstream functional consequences. Currently, both prospects remain challenging in part due to the technical complexity of analysis and lack of massively parallel analytical methods available. As such, there is a lack of functional annotations for most splice events compared to the well annotated functions of genes as described in Gene Ontology (Consortium et al., 2023). The consequences of splicing changes can be very different to the consequences of knockdown, underexpression or overexpression of the canonical isoform of a gene. For example, the two alternatively spliced protein isoforms of *BCL2*, Bcl-xL and Bcl-xS, have directly opposing effects on cell survival and apoptosis (Keller et al., 2023; Minnt et al., 1996). For other genes, alternative spliced non-coding isoforms themselves may mediate effects that are unrelated to the canonical protein isoform. For example, intron retention of steroid receptor RNA activator 1 (*SRA1*), results in a canonical non-coding transcript (*SRA1* RNA), as well as a transcript that produces a functional protein (SRA1P). While *SRA1* RNA acts as an RNA scaffold for the assembly of transcriptional co-factors to enhance transcription at estrogen receptor binding sites, SRA1P appears to have an antagonistic effect via an independent pathway (Dhamija and Menon, 2018; Hube et al., 2006). It is not known whether these are common modalities of functional regulation by alternative splicing as there could be many other examples which are yet to be discovered. Future functional studies could shed light on the extent to which effects are mediated by splicing and whether there are common modes of regulation by splicing.

There is the possibility that some splice events that are detectable have no functional significance – in particular, those arising from (unspliced) pre-mRNA. It has been shown that up to 25% (Gaidatzis et al., 2015) of total RNA reads can be attributed to unspliced RNA which could account for the improved correlation of splicing patterns with cell type due to the measurement of RNA velocity (La Manno et al., 2018). Nevertheless, we conject that as long as these are non-random and linked to the cell type, they may still be of meaning to cells and of use for sorting cell identity via data-driven methods, even though their origin and functional significance are unknown. It will thus be important to evaluate the reliability of using total RNA datasets in their entirety for data-driven analysis.

From an evolutionary standpoint, the difference in degeneracy factors for splicing and gene expression could be related to how gene regulatory mechanisms evolved, and how cell type diversity is created through alternative splicing. It has been speculated that alternative splicing provides a low energy pathway to transcript diversity and avoids changes which could be deleterious to an already well-optimised underlying genome sequence, or equally, diverse transcripts can act as a buffer to mask otherwise deleterious underlying genetic variants (Singh and Ahi, 2022; Verta and Jacobs, 2022). It thus remains to be seen if feature degeneracy of splicing and expression increases through evolutionary time, but such an analysis might prove challenging due to the requirement of obtaining high quality RNA to conduct analyses of a similar scale to the RNA Atlas.

## METHODS

### RNA Sequencing

The RNA-Seq data used for the RNA Atlas was obtained directly from the authors of the original RNA Atlas study but is also available in the National Center for Biotechnology Information’s Gene Expression Omnibus (GEO), accessible through GEO series accession number GSE138734. A description of the methods has been previously provided in the original RNA Atlas paper (Lorenzi et al., 2021). Briefly, library preparation, sequencing, and data processing of RNA-Seq was done for a total of 300 human samples were derived from 45 different tissue types, 162 primary cell types and 93 cell lines, of which 89 are cancer cell lines derived from 13 different types of cancer. RNA of primary cells and cancer cell lines was obtained from ScienCell Research Laboratories or isolated from samples collected at Ghent University Hospital. RNA samples from normal human tissues were obtained from Ambion and BioChain. Total RNA libraries were generated using the TruSeq Stranded Total RNA Library Prep Kit with Ribo-Zero Gold (Illumina) according to the manufacturer’s instructions using 1 μg of input RNA. PolyA capture RNA libraries were generated using the TruSeq Stranded mRNA Library Prep Kit (Illumina) according to the manufacturer’s instructions using 1 μg of input RNA. Library pools were quantified using the Standard Sensitivity NGS Fragment Kit on a Fragment Analyzer (Advanced Analytical). Pooled polyA and total RNA libraries were sequenced on a HiSeq 4000 instrument (Illumina) with paired-end 76 cycle reads.

### RNA-Seq data from the ENCODE project

We downloaded the call sets from the ENCODE portal (https://www.encodeproject.org/) (Luo et al., 2020) from all FASTQ files listed in the metadata file downloadable from the following query: https://www.encodeproject.org/metadata/?type=Experiment&control_type%21=%2A&status=release d&perturbed=false&assay_title=polyA+plus+RNA-seq&assay_title=total+RNA-seq&replicates.library.biosample.donor.organism.scientific_name=Homo+sapiens Experiments were filtered to only include those with more than 100 million paired-end reads, yielding 685 datasets.

### Reference genome alignment of RNA-Seq data

The quality of RNA-seq reads were confirmed with FastQC (version 0.11.9). Reads were aligned to the reference human genome (GRCh38 build) using STAR (version 2.7.2b) (Dobin et al., 2013). For alternative splicing analysis, paired FASTQ files were specified along with the following commands to perform two-pass mapping: `--outSAMtype BAM SortedByCoordinate--outSAMstrandfield intronMotif--outSJfilterReads Unique-- outFilterIntronMotifs RemoveNoncanonical--outFilterMultimapNmax 1-- twopassMode Basic`.

For gene expression analysis (RSEM only), paired FASTQ files were specified along with v98 of the Ensembl hg38 GTF file and the following commands to perform alignment to transcriptome:-- sjdbOverhang 99--quantMode TranscriptomeSAM.

### Gene expression analysis

For RSEM (version 1.3.3), reference sequences were prepared using the command `rsem-prepare-reference` using v98 of the hg38 assembly from Ensembl. Transcriptome-aligned BAM files from STAR were quantified using the command `rsem-calculate-expression` with the option `--paired-end`. For Salmon (version 1.10.1), a transcript index was built using the command `salmon index` with parameters `-t gencode`.`v38`.`transcripts`.`fa` and `-k 31`.

Paired FASTQ files were processed using `salmon quant-validateMappings` as per the mapping-based mode. TPM values were taken from the read counts of both tools; counts below 0.5 TPM were discarded.

Differential expression analysis was done using the limma package (version 3.56.2) in R with an FDR cutoff of 0.01 and a fold change cutoff of greater than 1.5.

### Alternative splicing analysis

For PSI-Sigma analysis (version 1.9k), a dummy comparison was set up by listing all BAM files into the document `groupa`.`txt` and `Total10_15601`.`Aligned`.`sortedByCoord`.`out`.`bam`, `Total10_15601-35019083`.`Aligned`.`sortedByCoord`.`out`.`bam` into the document `groupb`.`txt`. A universal database file was created by running the script `PSIsigma-db-v`.`1`.`0`.`pl` with parameters `Homo_sapiens`.`GRCh38`.`98`.`gtf 1 1 0`, and concatenating the resulting.tmp files to yield `psisigma_atlas_universal`.`db` and `psisigma_atlas_universal`.`bed`. The script `PSIsigma-ir-v`.`1`.`2`.`pl` was then run with parameters `psisigma_atlas_universal`.`bed ˂bamfile˃ 1`, with a separate instance for each BAM file. The script dummyai.pl was then run for each BAM file using the parameters `--gtf Homo_sapiens`.`GRCh38`.`98`.`gtf--name psisigma_atlas_universal--type 1--fmode 3--nread 1--irmode 0--denominator 1--adjp 0`. Counts below 5 reads were discarded. Differential splicing analysis was done using the package “DoubleExpSeq” (version 1.1) in R, with parameters shrink.method = “WEB” and fdr.level = 0.01. Splice events were considered differential if they also displayed a PSI change greater than 10% between comparisons.

For LeafCutter analysis (version 0.2.9), junction files were extracted from BAM files using the command `junctions extract` with parameters `-a 8-m 50-M 500000-s 1`. Intron clustering was performed using the script `leafcutter_cluster_regtools.py` with parameters `-m 50-l 500000`. Differential intron excision analysis was performed using the script `leafcutter_ds.R`.

### Sample clustering and dimensionality reduction

UMAP projections were generated using the umap package (version 0.2.10.0) in R. For clustering of splicing PSI and gene expression TPM profiles, an unsupervised fuzzy ensemble method was used. To do this, we used the som function in the kohonen package (version 3.0.11) in R, with parameters `somgrid(xdim =` ˂variable˃`, ydim =` ˂variable˃`, topo = “rectangular”, toroidal = FALSE), rlen = 100, dist`.`fcts = “sumofsquares”, mode = “online”`. Clustering was repeated with different xdim and ydim parameters such that 20 rectangular Kohonen self-organising maps were made, with dimensions 6 × 6 (xdim × ydim), 9 × 4, 19 × 2, 2 × 19, 5 × 8, 8 × 5, 6 × 7, 7 × 6, 4 × 11, 11 × 4, 23 × 2, 2 × 23, 6 × 8, 8 × 6, 2 × 25, 25 × 2, 2 × 26, 26 × 2, 6 × 9, 9 × 6. These dimensions were chosen to be uniformly distributed and centred around 45, the approximate number of clusters along the diagonal in the consensus matrix when sweeping across dimensions greater than 20. Clustering for all these dimensions was itself repeated with five different random number seeds.

Pairs of splice events or genes were considered to be “neighbours” if they co-clustered in more than 50% of the 100 maps generated. A clustering network was generated with each splice event or gene as a node, and the number of mutual neighbours as edge weightings. Fuzzy clusters were detected using ClusterONE in Cytoscape, with parameters: minimum cluster size = 25, node penalty = 2, haircut threshold = 0.1, merging method = “multi-pass”, similarity = “match coefficient”, overlap threshold = 0.8, seeding method = “from every node”. Clusters with a *P-*value greater than 0.05 were considered valid.

### Functional analysis of splice events

To obtain markers of differential splicing or expression for each cluster, comparisons were performed between the samples belonging to each ensemble cluster and all other the samples of the same category (i.e. tissue biopsy, primary cells, cell lines). This was done separately for each ensemble cluster in each dataset (total RNA and polyA capture datasets for PSI-Sigma, LeafCutter, RSEM and Salmon).

Gene ontology analyses were performed using the systemPipeR package (version 2.6.0) in R using the `GOHyperGAll` function. All differentially spliced or expressed genes in each ensemble cluster were enriched against a background of all known Ensembl genes (total RNA) or all Ensembl genes known to have a polyA tail (polyA capture).

To generate the annotated GO term network, the significantly enriched GO terms in totalRNA/PSI-Sigma and totalRNA/RSEM were summarised and overlaid onto a network representing the entire GO term parent-child hierarchy. The network view was generated in Cytoscape (version 3.10.1). Nodes were arranged using Prefuse Force Directed Layout based on the Jaccard index of the number of genes in common described by each GO term.

Summarisation of GO terms was necessary to generate a readable view of the GO term network whilst preserving the appearance of hotspots where GO terms were significantly enriched. To do this, a cost-optimisation algorithm was used to decide whether a more general parent term should replace a given GO term enriched in the dataset. For each dataset, the − log_10_ FDR values for each GO term were summed from all clusters into an aggregate logFDR (alFDR). A reward value was calculated for parent term(s) and compared to the alFDR of their child term using the following formula:

reward value = alFDR_parent_ − number of other child terms not enriched in dataset•*k* Where “other child terms” refer to the number of sub-terms linked to the parent term (regardless of whether they were enriched in the dataset), as defined by the GO term hierarchy.

If the reward value of the parent GO term is lower than the alFDR of the child term of interest, then the GO term of interest is not summarised. If the reward value of the parent GO term is higher than the alFDR of the child term of interest, then the parent GO term is taken, and the process is repeated for the previous parent as the new child term of interest. In this way, the algorithm traverses upwards in the GO term hierarchy until the reward value reaches a maximum or the highest level of the GO hierarchy is reached (Biological Process, Cellular Component or Molecular Function).

The algorithm balances the need to summarise similar GO terms against the loss of semantic specificity associated with taking a GO term at a higher level of hierarchy. If a parent is associated with a large number of child terms not enriched by the genes in the dataset, the reward function decreases as the parent term is more likely to describe biology that is not represented by the gene set. Adjustment of *k* changes how aggressive the summarising process is – we utilised a *k* value of 1. The nodes of the network diagram were the GO terms remaining after summarisation, coloured by the percentage of alFDR from splicing or expression.

### Sample classification

Cell classification was done using the scikit-learn 1.1.1 toolkit in Python 3.8.10.

Classifications were done for up to 18 test sizes for tissue biopsies, primary cells and cell lines: 5%, 10%, 15%, 20%, 25%, 30%, 35%, 40%, 45%, 50%, 55%, 60%, 65%, 70%, 75%, 80%, 85%, 90%.

Classifications at each of these test sizes were repeated 960 times. The following commands and parameters were used for each algorithm: `svm`.`SVC(kernel = “linear”, degree = 10) (svm)`, `LogisticRegression(solver = “liblinear”, max_iter = 1000) (logreg)`, `KNeighborsClassifier(n_neighbours = 2) (knn)`, `DecisionTreeClassifier(criterion = “entropy”) (decisiontree)`. Receiver-operator curves and associated AUROC values were generated using the pROC package (version 1.18.4) in R.

### Measuring feature degeneracy

Simple degeneracy to measure the presence or absence of a feature (splice event or gene) in samples was calculated the formula

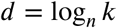

where *n* is the number of samples and *k* is the number of non-zero values.

Extended degeneracy to measure the proximity of feature values in a tolerance window was calculated using the same formula but using an updated value of *k*:

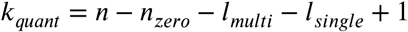

Where *n_zero_* is the number of zero values, *l_multi_* is the number of contiguous intervals created from the merging of more than one smaller interval, and *l_single_* is the number of unmerged intervals.

To generate *d*-Δ curves for each dataset, extended degeneracy values were calculated for each feature and then averaged. This was done for 1001 values of Δ:

{0, 0.001^4^, 0.002^4^, 0.003^4^,…, 0.998^4^, 0.999^4^, 1}. The 4^th^ power was taken because the steepest part of the *d*-Δ curves lay within the range from 10^−6^ to 10^−2^.

To generate the gradient *d*-Δ curves, the extended degeneracy value at every value of Δ was subtracted by that of its preceding value of Δ. This was done for all features, yielding a numerical matrix of gradient values. These values were clustered on a per feature basis using the som function in the kohonen package (version 3.0.11) in R, with parameters `somgrid(xdim = 100, ydim = 1, topo = “rectangular”, toroidal = FALSE, neighbourhood`.`fct = “gaussian”), rlen = 500, dist`.`fcts = “sumofsquares”, mode = “pbatch”`. Each cluster represented a bin, and gradient profiles from multiple features were averaged for each bin.

## ACKNOWLEDGEMENTS

A.L. acknowledges support from the Australian Government TRP scheme. M.R.W. acknowledges support from the Australian Research Council.

A.L. and M.R.W. thank Pieter Mestdagh and Ashwin Unnikrishnan for helpful discussions.

## REFERENCES

Ascensão-Ferreira, M., Martins-Silva, R., Saraiva-Agostinho, N. and Barbosa-Morais, N.L., 2024. betAS: intuitive analysis and visualisation of differential alternative splicing using beta distributions. RNA, 30, pp.337–353.

Boudreault, J., Wang, N., Ghozlan, M. and Lebrun, J.J., 2024. Transforming Growth Factor-β/Smad Signaling Inhibits Melanoma Cancer Stem Cell Self-Renewal, Tumor Formation and Metastasis. Cancers, 16(1), p.224.

Busch, A. and Hertel, K.J., 2015. Splicing predictions reliably classify different types of alternative splicing. RNA, 21, pp.813–823.

Cai, Q., He, B., Zhang, P., Zhao, Z., Peng, X., Zhang, Y., Xie, H. and Wang, X., 2020. Exploration of predictive and prognostic alternative splicing signatures in lung adenocarcinoma using machine learning methods. J Transl Med, 18, p.1–15.

Chen, M. and Manley, J.L., 2009. Mechanisms of alternative splicing regulation: insights from molecular and genomics approaches. Nat Rev Mol Cell Biol, 10, pp.741–754.

Consortium, T.G.O., Aleksander, S.A., Balhoff, J., Carbon, S., Cherry, J.M., Drabkin, H.J., Ebert, D., Feuermann, M., Gaudet, P., Harris, N.L. et al., 2023. The Gene Ontology knowledgebase in 2023. Genetics, 224, 224(1), p.iyad031.

Dam, S.H., Olsen, L.R. and Vitting-Seerup, K., 2023. Expression and splicing mediate distinct biological signals. BMC Biol, 21, p.1–10.

Dawicki-McKenna, J.M., Felix, A.J., Waxman, E.A., Cheng, C., Amado, D.A., Ranum, P.T., Bogush, A., Dungan, L.V., Maguire, J.A., Gagne, A.L., Heller, E.A., French, D.L., Davidson, B.L. and Prosser, B.L., 2023. Mapping PTBP2 binding in human brain identifies SYNGAP1 as a target for therapeutic splice switching. Nat Commun, 14(1), pp.1–20.

Dhamija, S. and Menon, M.B., 2018. Non-coding transcript variants of protein-coding genes – what are they good for? RNA Biol, 15, pp.1025–1031.

Dumitrascu, B., Villar, S., Mixon, D.G. and Engelhardt, B.E., 2021. Optimal marker gene selection for cell type discrimination in single cell analyses. Nat Commun, 12(1), pp.1–8.

Dvinge, H., 2018. Regulation of alternative mRNA splicing: old players and new perspectives. FEBS Lett, 592, pp.2987–3006.

Fanning, A.S. and Anderson, J.M., 2009. Zonula occludens-1 and-2 are cytosolic scaffolds that regulate the assembly of cellular junctions. Ann N Y Acad Sci, 1165(1), pp.113–120.

Feng, H., Moakley, D.F., Chen, S., McKenzie, M.G., Menon, V. and Zhang, C., 2021. Complexity and graded regulation of neuronal cell-type–specific alternative splicing revealed by single-cell RNA sequencing. Proc Natl Acad Sci U S A, 118, e2013056118.

Gaidatzis, D., Burger, L., Florescu, M. and Stadler, M.B., 2015. Analysis of intronic and exonic reads in RNA-seq data characterizes transcriptional and post-transcriptional regulation. Nat Biotechnol, 33(7), pp.722–729.

Ha, K.C.H., Sterne-Weiler, T., Morris, Q., Weatheritt, R.J. and Blencowe, B.J., 2021. Differential contribution of transcriptomic regulatory layers in the definition of neuronal identity. Nat Commun, 12(1), pp.1–12.

Hand, D.J. and Till, R.J., 2001. A simple generalisation of the area under the ROC curve for multiple class classification problems. Mach Learn, 45(2), pp.171–186.

Hube, F., Guo, J., Chooniedass-Kothari, S., Cooper, C., Hamedani, M.K., Dibrov, A.A., Blanchard, A.A.A., Wang, X., Deng, G., Myal, Y. and Leygue, E., 2006. Alternative splicing of the first intron of the steroid receptor RNA activator (SRA) participates in the generation of coding and noncoding RNA isoforms in breast cancer cell lines. DNA Cell Biol, 25(8), pp.418–428.

Johnson, N.T., Dhroso, A., Hughes, K.J. and Korkin, D., 2018. Biological classification with RNA-seq data: Can alternatively spliced transcript expression enhance Mach Learn classifiers?. RNA, 24(9), pp.1119–1132.

Katz, Y., Wang, E.T., Airoldi, E.M. and Burge, C.B., 2010. Analysis and design of RNA sequencing experiments for identifying isoform regulation. Nat Methods, 7(12), pp.1009–1015.

Keller, M.A., Huang, C.Y., Ivessa, A., Singh, S., Romanienko, P.J. and Nakamura, M., 2023. Bcl-x short-isoform is essential for maintaining homeostasis of multiple tissues. iScience, 26(4).

La Manno, G., Soldatov, R., Zeisel, A., Braun, E., Hochgerner, H., Petukhov, V., Lidschreiber, K., Kastriti, M.E., Lönnerberg, P., Furlan, A., Fan, J., Borm, L.E., Liu, Z., van Bruggen, D., Guo, J., He, X., Barker, R., Sundström, E., Castelo-Branco, G., Cramer, P., Adameyko, I., Linnarsson, S. and Kharchenko, P.V., 2018. RNA velocity of single cells. Nature, 560(7719), pp.494–498.

López-Martínez, A., Soblechero-Martín, P., De-La-puente-ovejero, L., Nogales-Gadea, G. and Arechavala-Gomeza, V., 2020. An overview of alternative splicing defects implicated in myotonic dystrophy type I. Genes, 11(10), p.1109.

Lorenzi, L., Chiu, H.S., Avila Cobos, F., Gross, S., Volders, P.J., Cannoodt, R., Nuytens, J., Vanderheyden, K., Anckaert, J., Lefever, S., Tay, A.P., de Bony, E.J., Trypsteen, W., Gysens, F., Vromman, M., Goovaerts, T., Hansen, T.B., Kuersten, S., Nijs, N., Taghon, T., Vermaelen, K., Bracke, K.R., Saeys, Y., De Meyer, T., Deshpande, N.P., Anande, G., Chen, T.W., Wilkins, M.R., Unnikrishnan, A., De Preter, K., Kjems, J., Koster, J., Schroth, G.P., Vandesompele, J., Sumazin, P. and Mestdagh, P., 2021. The RNA Atlas expands the catalog of human non-coding RNAs. Nat Biotechnol, 39(11), pp.1453–1465.

Luo, Y., Hitz, B.C., Gabdank, I., Hilton, J.A., Kagda, M.S., Lam, B., Myers, Z., Sud, P., Jou, J., Lin, K., Baymuradov, U.K., Graham, K., Litton, C., Miyasato, S.R., Strattan, J.S., Jolanki, O., Lee, J.W., Tanaka, F.Y., Adenekan, P., O’Neill, E. and Cherry, J.M., 2020. New developments on the Encyclopedia of DNA Elements (ENCODE) data portal. Nucleic Acids Res, 48(D1), pp.D882–D889.

Marasco, L.E. and Kornblihtt, A.R., 2022. The physiology of alternative splicing. Nat Rev Mol Cell Biol, 24(4), pp.242–254.

Matlin, A.J., Clark, F. and Smith, C.W.J., 2005. Understanding alternative splicing: towards a cellular code. Nat Rev Mol Cell Biol, 6(5), pp.386–398.

Mazin, P.V., Khaitovich, P., Cardoso-Moreira, M. and Kaessmann, H., 2021. Alternative splicing during mammalian organ development. Nature Genet, 53(6), pp.925–934.

Minnt, A.J., Boise, L.H. and Thompson, C.B., 1996. Bcl-XS antagonizes the protective effects of Bcl-xL. J Biol Chem, 271(10), pp.6306–6312.

Najar, C.F.B.A., Yosef, N. and Lareau, L.F., 2020. Coverage-dependent bias creates the appearance of binary splicing in single cells. eLife, 9, p.e54603.

Park, J.W., Fu, S., Huang, B. and Xu, R.H., 2020. Alternative splicing in mesenchymal stem cell differentiation. Stem Cells, 38, pp.1229–1240.

Pickrell, J.K., Pai, A.A., Gilad, Y. and Pritchard, J.K., 2010. Noisy splicing drives mRNA isoform diversity in human cells. PLoS Genet, 6, pp.1–11.

Pullin, J.M. and McCarthy, D.J., 2024. A comparison of marker gene selection methods for single-cell RNA sequencing data. Genome Biol, 25, pp.1–37.

Reixachs-Solé, M. and Eyras, E., 2022. Uncovering the impacts of alternative splicing on the proteome with current omics techniques. Wiley Interdiscip Rev RNA, 13, p.e1707.

Shalek, A.K., Satija, R., Adiconis, X., Gertner, R.S., Gaublomme, J.T., Raychowdhury, R., Schwartz, S., Yosef, N., Malboeuf, C., Lu, D., Trombetta, J.J., Gennert, D., Gnirke, A., Goren, A., Hacohen, N., Levin, J.Z., Park, H. and Regev, A., 2013. Single-cell transcriptomics reveals bimodality in expression and splicing in immune cells. Nature, 498(7453), pp.236–240.

Singh, P. and Ahi, E.P., 2022. The importance of alternative splicing in adaptive evolution. Mol Ecol, 31, pp.1928–1938.

Slaff, B., Radens, C.M., Jewell, P., Jha, A., Lahens, N.F., Grant, G.R., Thomas-Tikhonenko, A., Lynch, K.W. and Barash, Y., 2021. MOCCASIN: a method for correcting for known and unknown confounders in RNA splicing analysis. Nat Commun, 12(1), p.1–9.

Stamm, S., Ben-Ari, S., Rafalska, I., Tang, Y., Zhang, Z., Toiber, D., Thanaraj, T.A. and Soreq, H., 2005. Function of alternative splicing. Gene, 344, pp.1–20.

Tapial, J., Ha, K.C.H., Sterne-Weiler, T., Gohr, A., Braunschweig, U., Hermoso-Pulido, A., Quesnel-Vallières, M., Permanyer, J., Sodaei, R., Marquez, Y., Cozzuto, L., Wang, X., Gómez-Velázquez, M., Rayon, T., Manzanares, M., Ponomarenko, J., Blencowe, B.J. and Irimia, M., 2017. An atlas of alternative splicing profiles and functional associations reveals new regulatory programs and genes that simultaneously express multiple major isoforms. Genome Res, 27, pp.1759–1768.

Verta, J.P. and Jacobs, A., 2022. The role of alternative splicing in adaptation and evolution. Trends Ecol Evol, 37, pp.299–308.

Wan, Y. and Larson, D.R., 2018. Splicing heterogeneity: separating signal from noise. Genome Biol, 19(1), pp.1–10.

Wang, E.T., Sandberg, R., Luo, S., Khrebtukova, I., Zhang, L., Mayr, C., Kingsmore, S.F., Schroth, G.P. and Burge, C.B., 2008. Alternative isoform regulation in human tissue transcriptomes. Nature, 456(7221), pp.470–476.

Wright, C.J., Smith, C.W.J. and Jiggins, C.D., 2022. Alternative splicing as a source of phenotypic diversity. Nat Rev Genet, 23(11), pp.697–710.

Xiao, Y., Gao, L., Zhao, X., Zhao, W., Mai, L., Ma, C., Han, Y. and Li, X., 2024. Novel prognostic alternative splicing events in colorectal cancer: Impact on immune infiltration and therapy response. Int Immunopharmacol, 139, p.112603.

Zeng, H., 2022. What is a cell type and how to define it? Cell, 185, pp.2739–2755.

Zhang, C., Frias, M.A., Mele, A., Ruggiu, M., Eom, T., Marney, C.B., Wang, H., Licatalosi, D.D., Fak, J.J. and Darnell, R.B., 2010. Integrative modeling defines the nova splicing-regulatory network and its combinatorial controls. Science, 329, pp.439–443.

Zhang, Y., Qian, J., Gu, C. and Yang, Y., 2021. Alternative splicing and cancer: a systematic review. Signal Transduct Target Ther, 6(1), pp.1–14.

Zhao, H., 2012. Membrane trafficking in osteoblasts and osteoclasts: New avenues for understanding and treating skeletal diseases. Traffic, 13, pp.1307–1314.

Zhao, L., Wang, J., Li, Y., Song, T., Wu, Y., Fang, S., Bu, D., Li, H., Sun, L., Pei, D., Zheng, Y., Huang, J., Xu, M., Chen, R., Zhao, Y. and He, S., 2021. NONCODEV6: an updated database dedicated to long non-coding RNA annotation in both animals and plants. Nucleic Acids Res, 49, pp.D165–D171.

Zhou, J., Zhao, S. and Dunker, A.K., 2018. Intrinsically disordered proteins link alternative splicing and post-translational modifications to complex cell signaling and regulation. J Mol Biol, 430, pp.2342–2359.

Zhou, X., Zhang, Z., Feng, J.Q., Dusevich, V.M., Sinha, K., Zhang, H., Darnay, B.G. and De Crombrugghe, B., 2010. Multiple functions of Osterix are required for bone growth and homeostasis in postnatal mice. Proc Natl Acad Sci U S A, 107, pp.12919–12924.

Zhou, X.G., Ren, L.F., Li, Y.T., Zhang, M., Yu, Y.D. and Yu, J., 2010. The next-generation sequencing technology: A technology review and future perspective. Sci China Life Sci, 53(1), pp.44–57.

Zhou, Z., Qu, J., He, L., Peng, H., Chen, P. and Zhou, Y., 2018. α6-Integrin alternative splicing: distinct cytoplasmic variants in stem cell fate specification and niche interaction. Stem Cell Res Ther, 9, pp.1–7.

